# Forecasting autism gene discovery with machine learning and genome-scale data

**DOI:** 10.1101/370601

**Authors:** Leo Brueggeman, Tanner Koomar, Jacob J Michaelson

## Abstract

**Background:** Genes are one of the most powerful windows into the biology of autism, and it has been estimated that perhaps a thousand or more genes may confer risk. However, less than 100 genes are currently viewed as having robust enough evidence to be considered true "autism genes". Massive genetic studies are underway to produce data to implicate additional genes, but this approach, although necessary, is costly and slow-moving.

**Methods:** We approach autism gene discovery as a machine learning problem, rather than a genetic association problem, and use genome-scale data as predictors for identifying further genes that have similar properties in the feature space compared to established autism risk genes. This approach, which we call forecASD, integrates spatiotemporal gene expression, heterogeneous network data, and previous gene-level predictors of autism association into an ensemble classifier that yields a single score that indexes each gene’s evidence for being involved in the etiology of autism.

**Results:** We demonstrate that forecASD has substantially increased sensitivity and specificity compared to previous gene-level predictors of autism association, including genetic measures such as TADA. On an independent test set, consisting of newly-released pilot data from the SPARK Genomics Consortium, we show that forecASD best predicts which genes will have an excess of likely gene disrupting (LGD) *de novo* mutations. We further use independent data from a recent post mortem study of case/control gene expression to show that forecASD is also a significant predictor of genes implicated in ASD through differential expression. Using forecASD results, we show which molecular pathways are currently under-represented in the autism literature and likely represent under-appreciated biological mechanisms of autism. Finally, forecASD correctly predicted 12 of 16 genes implicated at FDR=0.2 by the latest ASD gene discovery study, while also identifying the most likely false positives among the candidate genes.

**Conclusions:** These results demonstrate that forecASD bridges the gap between genetic- and expression-based ASD gene discovery, and provides a data-driven replacement to much of the manual filtering and curation that is a critical step in ensuring the robustness of gene discovery studies.

## Background

Autism Spectrum Disorder (ASD) is a heterogeneous grouping of developmental disorders caused by a range of genetic and environmental factors. The core diagnostic features of ASD, which manifest at a young age, are impairments in social communication and restrictive and repetitive behaviors and interests. Evidence for the role of genetics in ASD is strong, with monozygotic twins having near 90% concordance of ASD diagnosis(1). Further population and twin studies have confirmed these findings(2), and further estimated the narrow-sense heritability of ASD to be in the range of 50-95%.

While there is an abundance of evidence for the role of genetics in autism, our understanding of the genetic etiology of the disorder is still limited. It is estimated that there may over 1000 genes which contribute to autism risk(3). However, the current list of high-confidence autism genes stands at 84 genes(4). This discrepancy is partly explained by the relatively limited number of genomic studies compared with the vast genetic heterogeneity underlying autism.

To close this gap between the number of anticipated and known autism genes, several network-biology approaches have been applied in the past decade. These studies leverage large, publicly-available datasets to add context and amplify the genetic signals observed through sequencing studies. These network-biology studies have predicted genes that then became bona fide autism genes(5), but have fallen short of providing a useful genome-wide metric that indicates the evidence of autism involvement for every gene. More recently, machine learning based methods have used gene interaction networks(6) and cell-specific expression profiles(7) to predict gene involvement in autism. Importantly, the results of these studies lead to a quantitative metric that scores every gene in the genome according to evidence of a role in autism. Despite the demonstrated effectiveness of these studies in prioritizing autism risk genes, our preliminary investigations suggested there was still room for appreciable improvement in the form of the classification algorithm, the training set, and the predictors used. In particular, these approaches do not incorporate indicators of autism involvement that are based on genetic association (e.g., TADA scores) into their predictive features.

We introduce a new score, forecASD, that integrates prior network-biology approaches, scores of genetic association, brain gene expression, and topological information from large gene interaction networks relevant to the brain into a single gene-level score for autism involvement. We show that forecASD successfully outperforms existing methods in a diverse range of gene and mutation prioritization tasks. Further, using the recent sequencing studies MSSNG(8) and SPARK(9), we show that forecASD generalizes to previously unseen data. Importantly, this generalization holds even when excluding genes with known links to autism, emphasizing forecASD’s ability to identify novel ASD genes. We also demonstrate that forecASD correctly predicts 12 of 16 genes implicated at FDR=0.2 by the latest ASD gene discovery study, while also identifying the most likely false positives among the candidate genes. Through comparing the top decile of forecASD identified genes (1787 genes; hereafter forecASD genes) with known autism genes, we identify numerous biological pathways that are currently underrepresented in our understanding of autism risk. By reanalyzing the results of autism brain differential gene expression studies, we show that the current list of known autism genes is significantly depleted for upregulated biological pathways, whereas forecASD captures both up- and downregulated pathways. We show that direction of differential expression is related to haploinsufficiency status, with low pLI genes showing a trend towards upregulation. Importantly, this relationship between direction of differential expression and pLI is dependent on forecASD gene inclusion, signifying forecASD’s ability to capture low- and high-pLI disease genes. Through these studies, we show evidence that current methods of autism gene discovery have biases, and that forecASD mitigates these biases through its integrative approach, thus providing a view of the full spectrum of genes and biological pathways underlying autism.

## Methods

### Overview

The forecASD method relies upon stacked Random Forest models, organized in two levels (shown in Figure 1). In the first level, two models are trained using BrainSpan(10) gene expression and the STRING(11) shortest paths network as features, respectively. Our training dataset consists of high-confidence genes scored in SFARI gene(4) as either 1 or 2 (SFARI HC genes), and 1,000 random background genes not contained within SFARI gene. These two models produce genome-wide predictions for autism involvement. These scores are then used as features in the second level’s Random Forest model, along with other genome-wide scores obtained from previous studies.

**Figure 1.**
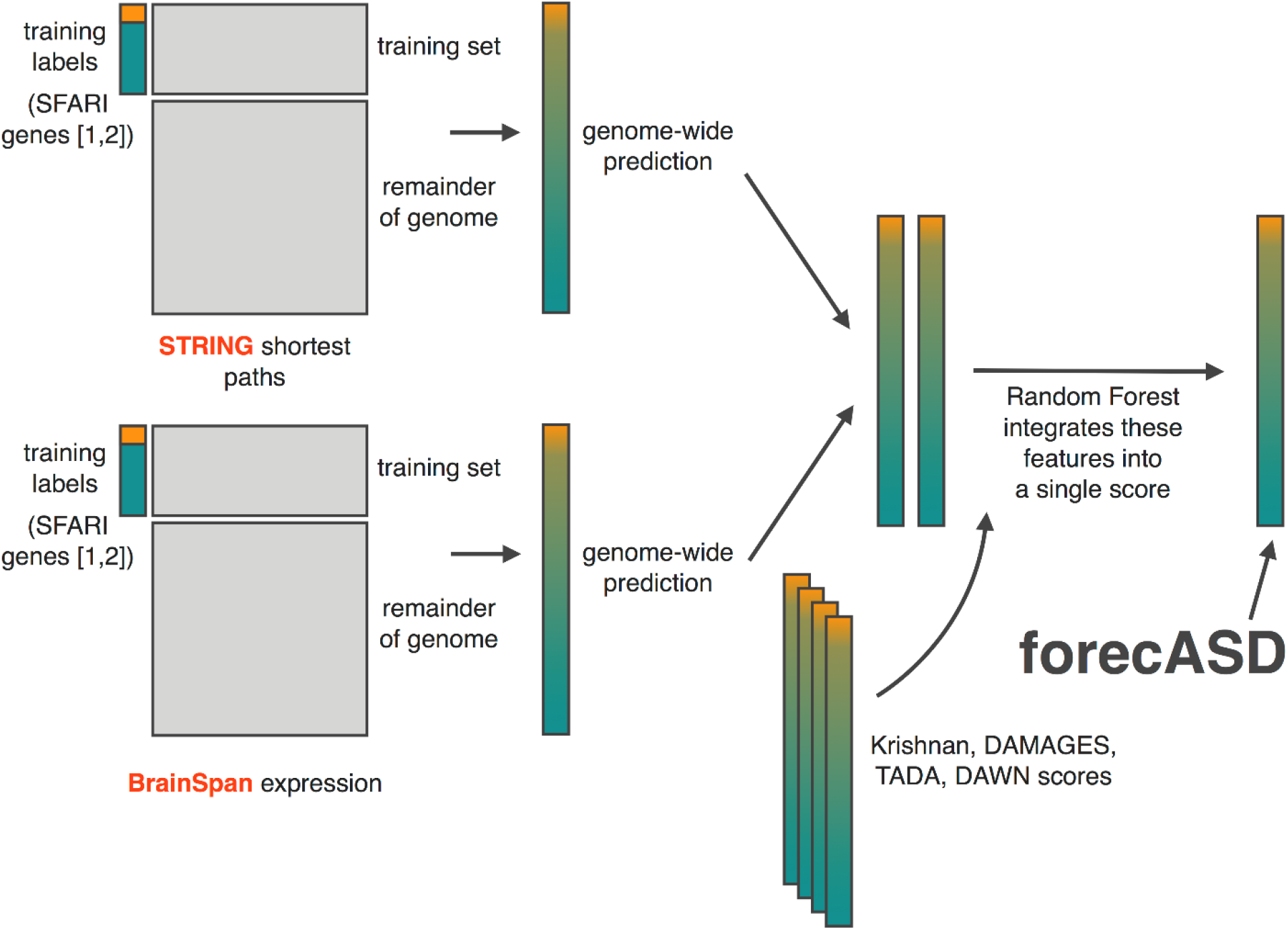
Overview of forecASD. Two Random Forest classifiers, one using BrainSpan gene expression and the other using the STRING network as predictors, are trained to discriminate high confidence autism genes (SFARI HC, scores 1 and 2) from a set of 1,000 genes drawn randomly from those not listed at all in the SFARI Gene database. Predictions are then made on the remainder of the genome, and these are combined with the out-of-bag (OOB) estimates from the training process to yield a prediction for each gene in the genome. A subsequent classifier is then trained using the output of these two RFs and previously published autism gene scores as predictive features, and again predictions are made on the remainder of the genome, with OOB predictions being used for those genes in the training set. The RF vote proportion for class “autism gene” is then the final forecASD score.

### BrainSpan, STRING, and TADA data assembly

BrainSpan data was obtained from the Allen Institute, and brain regions containing fewer than 20 samples were excluded. This filtered BrainSpan dataset was loess-smoothed, with the purpose of reducing noise and imputing missing data points.

The STRING database(11) was thresholded at their recommended score of 0.4, and transformed into a gene by gene matrix with each cell representing the shortest path between two genes.

TADA summary statistics were downloaded from the largest meta-analysis for autism available at the time of publication*(12)*. TADA summary statistics were also obtained from the secondary supplementary table of another comprehensive study of autism risk*(3)*. All available TADA summary statistics were used as features in the final model, with tadaFdrAscSscExomeSscAgpSmallDel*(12)* used as a representative comparator in the ROC curve displayed in Figure 3.

### Model training and genome wide prediction

We used a stacked Random Forest classifier to generate genome-wide predictions of autism gene involvement. All models were trained using SFARI HC genes as positive examples (of which there are 76 common to both STRING & BrainSpan), and a randomly sampled set of 1,000 background genes (i.e., not listed in the SFARI Gene database) as negative examples.

The first level of our stacked model consists of two genome-wide scores based on data from BrainSpan or the STRING interaction network. The features used in training these two models include the loess-smoothed observations in the BrainSpan database, and the STRING shortest path matrix, respectively. The random forest models were trained with 1000 total trees constructed, and the strata option enabled to insure a balance of 70 positive and 70 negative training examples during the construction of each tree. Given the large number of features for the STRING-based random forest model, we performed feature selection, wherein each feature not used in any of the constructed trees was dropped. This variable selection step was repeated until the final model contained only features which were selected at least once during tree construction. With the STRING and BrainSpan models, we then predicted autism involvement scores for the remaining genes not included in our training set. These scores are in Supplementary Table 1 in the columns BrainSpan_score and STRING_score. Scores for training set genes are the out-of-bag estimates.

We used these scores, along with DAWN(5), TADA(12)^,(3)^, DAMAGES(7), and the score from Krishnan *et al.(6)* score, as predictive features in a final Random Forest, using the same training labels described previously. Genome-wide predictions were then obtained, again using out-of-bag estimates for training set genes. This final score is listed under forecASD_score in Supplementary Table 1.

### SPARK and MSSNG data sources

De novo mutation (DNM) data from the MSSNG dataset was obtained through the *de novo db* database(6). Mutations were filtered for LGD or missense status. De novo mutation data was obtained from the SPARK dataset from the consortium’s recently released de novo mutation table. For both SPARK and MSSNG, only DNMs for probands were used.

### Pathway enrichment and comparison with case/control brain gene expression data

We used Reactome annotations(13), and unless otherwise noted, PantherDB(14) to assess functional enrichment in both forecASD genes and SFARI HC genes using Fisher’s method. Odds ratios and p-values were used to compare these two prioritization methods (Fig. 4) in terms of the pathways they implicate. The full list of results of these enrichment analyses are provided in Supplemental Table 2. Statistical analyses described in results and discussion were all performed in R(15) using either glm() or fisher.test(). Pathway-summarized haploinsufficiency (pLI: probability of loss-of-function intolerance(16)) was calculated by counting the proportion of genes in a Reactome pathway satisfying pLI>0.9. Gene-wise and pathway-level comparisons with ASD case/control brain gene expression data were performed using frontal cortex RNA-seq summary statistics from Gandal *et al.(17)*. Our preliminary tests showed that both SFARI HC and forecASD showed the highest agreement with expression data from the frontal cortex.

### Class and functional enrichment of top forecASD genes

Data used for functional enrichment in Figure 2D was taken from PubMed, STRING(11), and BrainSpan(10), using forecASD genes as the subject. PubMed literature enrichment scores were calculated by summing total mentions of the gene list in abstracts also containing the word autism. The network interaction scores were derived using the STRING database, accessed via the STRINGdb package(18) in R(15). Using a score threshold of 0.4, we keep all STRING interactions between top forecASD genes. The total number of interactions above this threshold is then summed. Fetal brain coexpression scores are based on average Pearson correlation between top scoring forecASD genes in early developmental timepoints in the BrainSpan dataset. Given these three functional enrichment scores, average background values were permuted by randomly drawing a set of 1787 genes 1000 times. P-values and enrichment were computed relative to the permuted samples. Datasets used in the class enrichment in Figure 2D were taken from Sugathan *et al.(19)*, Darnel *et al.(20)*, and Abrahams *et al.(4)*. P-values were computed by the hypergeometric statistical test of overlap between forecASD genes and these three gene sets.

**Figure 2.**
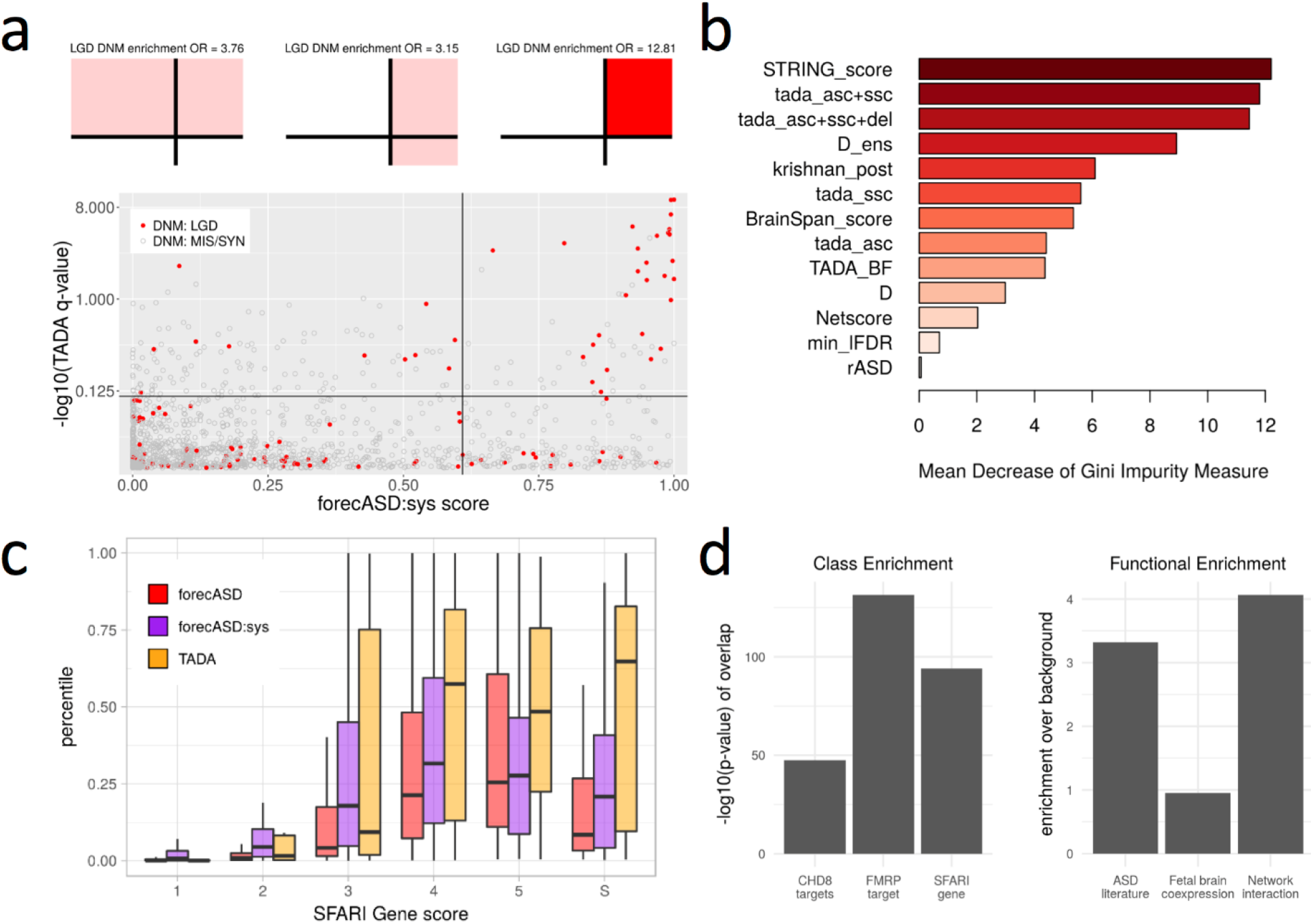
Prioritization of de novo likely gene-disrupting mutations and enrichment of gene sets in forecASD. Training a limited model, forecASD:sys, using brain gene expression and interaction data shows optimal prioritization of de novo LGDs when combined with a genetic measure of autism association (a). Building the full forecASD model, we test all features for their informativeness, finding that the STRING score is primary (b). Using the three mentioned scores, we assess their genome-wide ranking of SFARI genes at all levels, and find that the full forecASD model at least ties, and often significantly outperforms TADA and forecASD:sys in the prioritization of SFARI genes (c). As an initial assessment of forecASD prioritized genes, we find the top decile of genes ranked by forecASD (1787 genes) shows enrichment typical of classical autism genes (d).

### Cluster analysis of top scoring forecASD genes

Using the STRING database, interactions were obtained for forecASD genes and loaded into a network using the igraph package(21) in R(15). No filter for interaction strength was enforced. Hierarchical greedy clustering based on optimization of the modularity score(22) was performed using the fastgreedy.community function in the igraph package. Clustering was performed iteratively, with clusters containing more than 200 genes being subject to further clustering. Clusters with fewer than 30 genes were discarded. The annotated network of clusters was loaded into Cytoscape(23), using the STRING application(18). Functional enrichment of clusters was assessed using the STRING application in Cytoscape, with the p-value threshold set to 0.05. For annotation of the network plot shown in figure 6A, either the top annotated term or commonality between several top terms was chosen as representative. The p-value of enrichment with pLI scores was performed using Fisher’s exact test of genes within each cluster with a pLI score above 0.5. The p-value for overlap with SFARI HC genes was performed using the hypergeometric test, assuming a background of 18,000 total genes.

## Results

### forecASD model and performance

The goal of our approach was to create a gene-wise score that indexes the level of evidence for involvement in ASD using both systems biology (i.e., network and transcriptional data) and genetic features. An initial forecASD systems biology model was built (forecASD:sys) using only BrainSpan expression and the STRING database shortest paths matrices as features. This model was trained on the high confidence set of 76 SFARI genes scoring 1 or 2 (SFARI HC genes), with negative training labels assigned to 1,000 background genes that were not listed in the SFARI gene database.

As an initial test of performance, we scored genes hit by coding *de novo* mutations (DNMs) in the recently published MSSNG study. As shown in Figure 2A, there is a significant enrichment of likely gene disrupting (LGD) DNMs in the 90^th^ percentile of both the TADA p-value (OR = 3.76, P = 5.45 × 10-9) and the forecASD:sys scores (OR = 3.15, P = 7.33 × 10^-8^). However, by far the greatest enrichment (OR = 12.81, P < 2.2 × 10^-16^) is seen when restricting to DNMs passing both a TADA q-value and forecASD:sys 90^th^ percentile threshold.

To leverage both the genetic signal and the systems biology signal, we next built the final forecASD model, which incorporates forecASD:sys, the Krishnan *et al.* score(6), DAMAGES(7), DAWN(5), and several TADA genetic scores from two recent studies(3)^,(12)^. After training the forecASD model, we visualized the variable importance in figure 2B by mean decrease in the Gini impurity measure. The most informative feature was the STRING score from the forecASD:sys model, followed closely by two TADA score variables.

To facilitate a comparison with manually curated gene prioritizations, we scored all genes in the SFARI gene database using forecASD, forecASD:sys, and the most comprehensive TADA feature in the forecASD model. Shown in figure 2C, the forecASD model ranks SFARI genes scoring 3, 4, 5 and syndromic-only as significantly more autism-related than TADA (P: 7.7×10^-4^, 4.7×10^-11^, 2.3×10^-4^, 7.7×10^-6^). The forecASD model also significantly outperforms the limited forecASD:sys model in gene categories 2, 3, and 4 (P: 8.4×10^-5^, 2.15×10^-7^, 4.0×10^-5^, respectively). In all cases, forecASD prioritizes SFARI genes as well, or better than TADA and forecASD:sys.

As an initial validation of genes prioritized by forecASD, we tested for an enrichment of gene sets and characteristics well known to be overrepresented in autism genes (Fig. 2D). We first performed several overrepresentation tests and found that genes receiving forecASD scores in the top decile (1,787 genes, referred to as forecASD genes) had a significant overlap with known targets of CHD8 (P < 1 × 10^-16^), FMRP (P < 1 × 10^-16^), and the full SFARI gene database (P < 1 × 10^-16^). We next performed a series of functional enrichment tests, comparing forecASD genes to randomly sampled sets of background genes. Text mining in PubMed showed that forecASD genes were significantly overrepresented in abstracts which mention autism (P < 0.001). Given the established role of autism genes early in fetal development, we next tested and found that forecASD genes showed significantly higher rates of coexpression across all regions of the fetal brain (P < 0.001). Lastly, forecASD genes were shown to have significantly enriched rates of interaction in the STRING database (P < 0.001).

We next tested the ability of these scores to discriminate both high confidence (Fig. 3A) and trending (Fig. 3B) autism genes from negative background genes. High confidence autism genes (SFARI HC) are defined as scoring 1 or 2 in SFARI Gene, with trending autism genes scoring 3. Importantly, the negative set of non-autism genes was sampled to have the same background mutation rate as the autism genes (P>0.1 by the Kolmogorov-Smirnov test). In both comparisons, forecASD showed the highest level of performance of all methods tested (AUC=0.97 for SFARI 1+2 and AUC=0.82 for SFARI Gene score 3; Fig. 3). Furthermore, while the SFARI HC genes were used to train the forecASD model, only “out of bag” predictions were used as the forecASD score for those genes, i.e., only those trees where the gene was not included in the bootstrap sample voted for the class of the gene. None of the trending autism genes (Fig. 3B) were used to train forecASD, and consequently they provide an unbiased estimate of performance.

**Figure 3.**
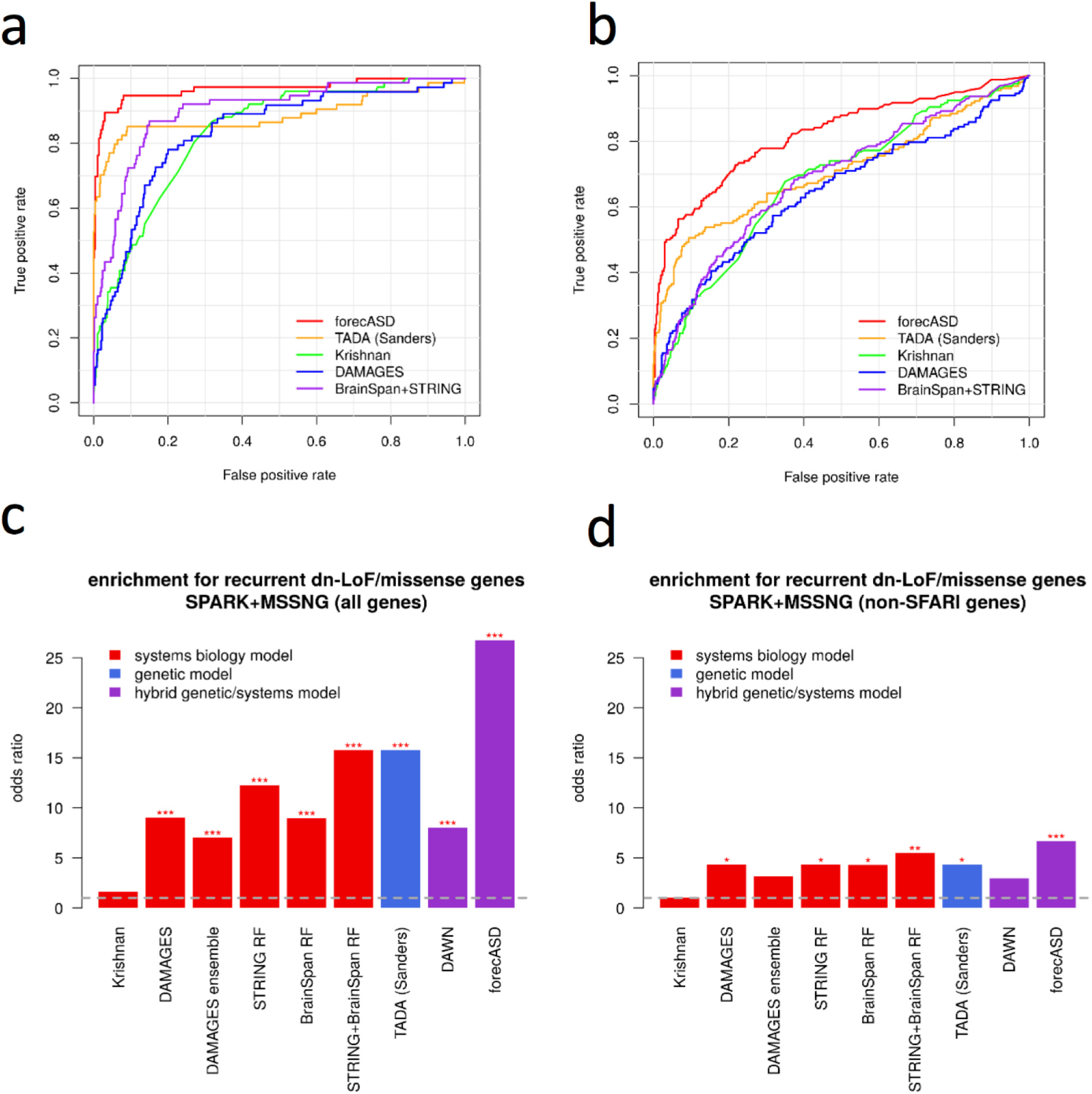
Comparison of forecASD with prior models of autism gene prioritization. To compare forecASD with competitors, we evaluate performance by each methods’ ability to prioritize SFARI genes and genes which were subject to recurrent de novo loss-of-function or missense mutations. Starting with SFARI genes scoring 1 or 2 as a positive set and size-matched random background genes as the negative set, forecASD out-of-bag estimates showed superior classification over all methods (a). In a fully unbiased test, forecASD estimates also showed superior classification of trending SFARI genes (score: 3) over all other methods (b). Using two sequencing cohorts which no methods draw information from, the top decile of forecASD genes (1787 genes) shows the greatest overlap with genes containing recurrent de novo loss-of-function and missense mutations (c). When excluding genes in the SFARI gene database, forecASD still shows superior prioritization of genes accumulating de novo mutations (d).

### Generalization to new data: *de novo* mutation enrichment

To compare forecASD and prior methods’ ability to generalize to new data, we combined two recently released autism genetics resources. Specifically, we used *de novo* mutations in gene regions from the SPARK(9) and MSSNG(8) cohorts. Importantly, none of our model training used information from these studies, thus any subsequent validation is unbiased.

We first compared forecASD and competing ASD gene scores with respect to enrichment of genes with recurrent *de novo* loss of function and damaging missense mutations in probands. forecASD significantly outperformed all prior approaches (OR=26.8, P=3.1×10^-24^; Fig. 3C). We next tested whether forecASD continued to show significant enrichment when known autism genes (here, any gene listed in the SFARI gene database, regardless of score) were excluded (Fig. 3D), since the ideal method should detect both known and potentially novel autism genes. forecASD had superior performance in this test as well (OR=6.7, P=0.0004), with most of the other external methods lacking a statistically significant enrichment over baseline.

### Functional enrichment and clustering of forecASD genes

Having demonstrated the predictive performance characteristics of forecASD, we next turned to practical applications that could further illuminate the underlying biological mechanisms at play in autism. Functional enrichment using Reactome annotations showed that forecASD genes are highly enriched for pathways known to play an important role in autism etiology, including chromatin modification, synaptic transmission, and developmental biology (full list in Supplemental Table 2). To highlight new biological themes that forecASD detects but that are not clear from the list of SFARI HC genes, we prioritized pathways based on differential enrichment (Fig. 4). Figure 4A highlights pathways that were represented in SFARI HC genes, but that showed significantly greater enrichment in forecASD genes. Figure 4B shows a sampling of the most significant forecASD pathways not represented by any SFARI HC gene, thus highlighting under-appreciated mechanisms in autism.

**Figure 4.**
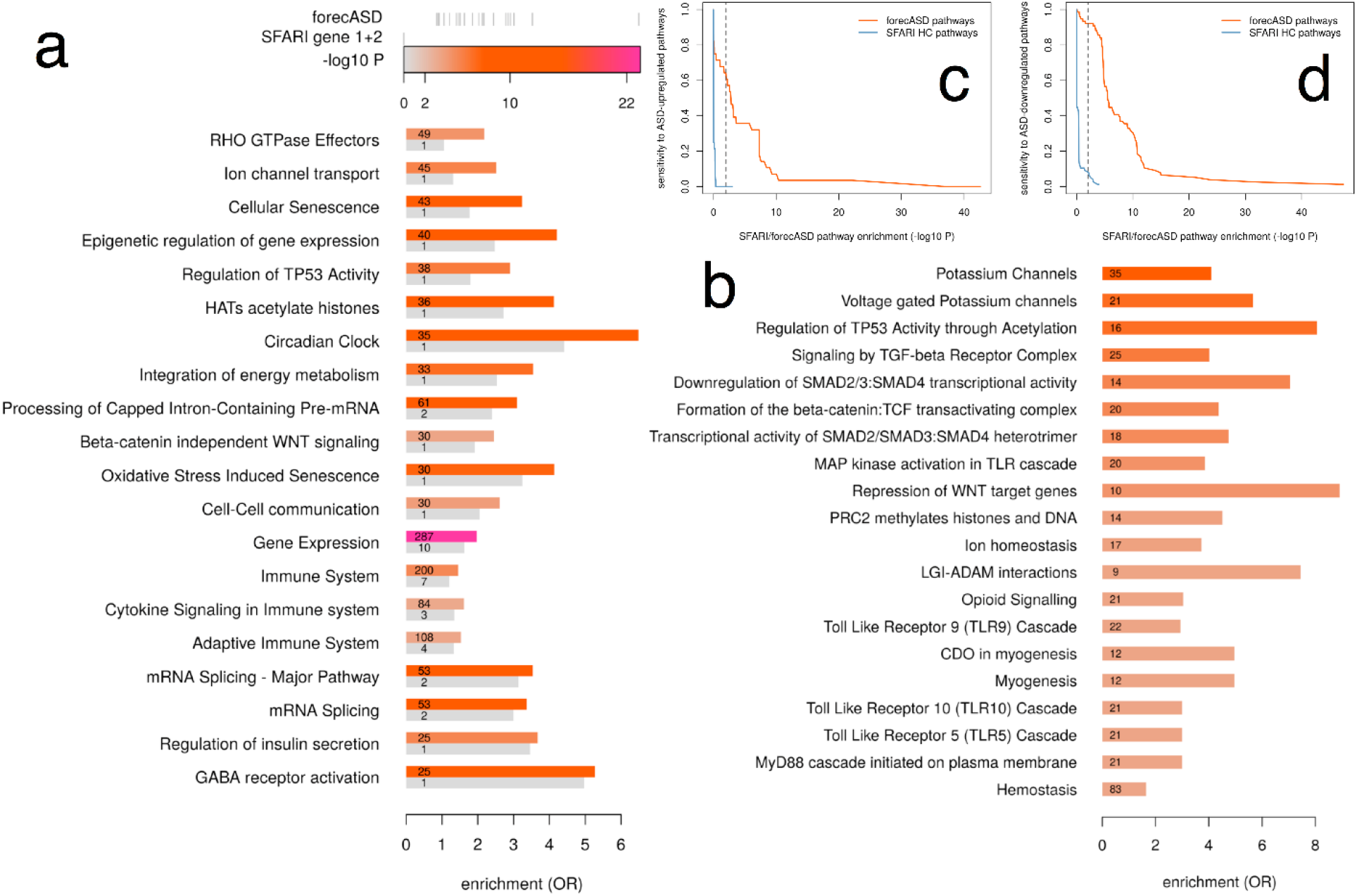
forecASD-specific pathway enrichment and sensitivity to gene expression-implicated pathways. When testing the top-decile genes according to forecASD for Reactome pathway enrichment, pathways emerged that were represented, but not enriched in the SFARI HC list (a). Other pathways were highly enriched in forecASD genes that were not represented at all in the SFARI HC list, even though they have associated literature suggesting a role in autism (b). forecASD is more sensitive than SFARI HC to pathways that are differentially regulated in the brains of individuals with autism, particularly in ASD-upregulated pathways (c), but also in downregulated pathways (d). Using the top decile of TADA −log10 FDR genes showed similar sensitivity to SFARI HC (not shown), suggesting that rare variant approaches may be less sensitive in implicating genes found through gene expression studies.

While SFARI HC genes show a strong bias toward genes with high pLI (P<0.001, Fisher’s exact test; Fig. 5A), forecASD is significantly less biased (P<0.001, Fisher’s exact test). We also discovered a significant relationship between pLI and differential expression (DE) t-statistics in case/control brain gene expression studies(17) (beta=-0.13, t-statistic-4.3, P=1.9×10^-5^, Fig. 5B), potentially exposing a form of bias in current gene discovery approaches that leads to under-ascertainment of ASD risk genes with low pLI and upregulation in ASD cases. We also found a significant interaction between forecASD and pLI (*F*=54.1, P=3.9×10^-24^) such that the pLI-expression relationship exists among forecASD genes (beta=-0.24, t=-2.6, P=0.009; Fig. 5D) but is absent in non-forecASD genes (beta=0.004, t=0.1, P=0.91; Fig. 5C).

**Figure 5.**
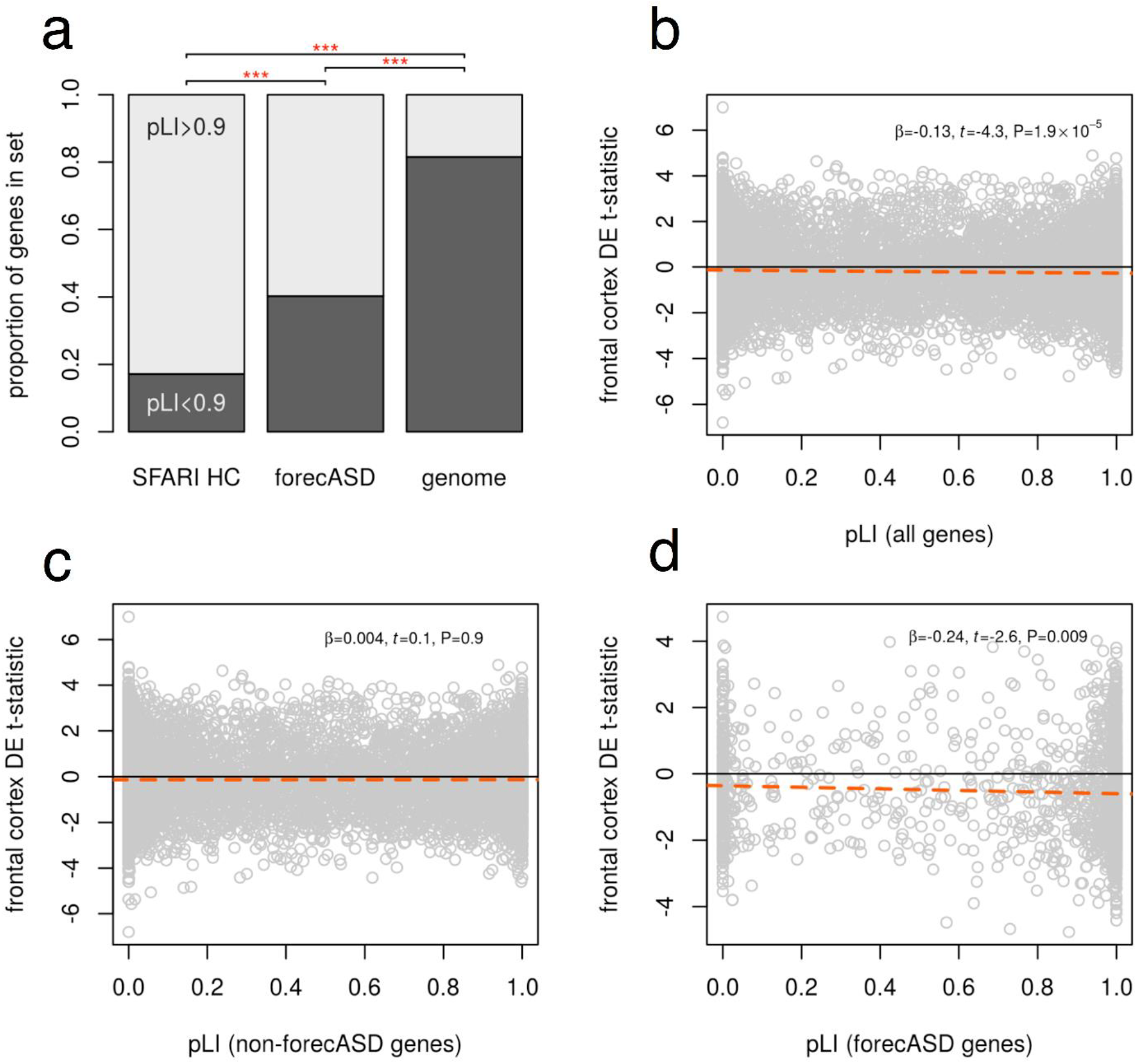
Relationship between pLI and ASD-specific up- and down-regulation of brain gene expression. SFARI HC is strongly biased toward genes with high pLI (a), while forecASD is significantly less biased. We found a significant relationship between pLI and differential expression (DE) in the brains of autism cases (b), such that low pLI genes tend toward upregulation in cases, while high pLI genes tend to be downregulated. We also observed a significant interaction between forecASD and pLI such that the observed pLI-DE trend (b) is absent in non-forecASD genes (c), and present and significant among forecASD genes (d). We propose that the presence of the pLI-DE trend is a hallmark of ASD risk genes, and an optimal ASD gene prioritization method will concentrate the trend among risk genes and remove it from non-risk genes. Notably, no threshold of TADA (tested to the 50^th^ percentile) was able to remove the trend from the non-prioritized genes, suggesting the persistence of residual risk genes that were not selected.

**Figure 6.**
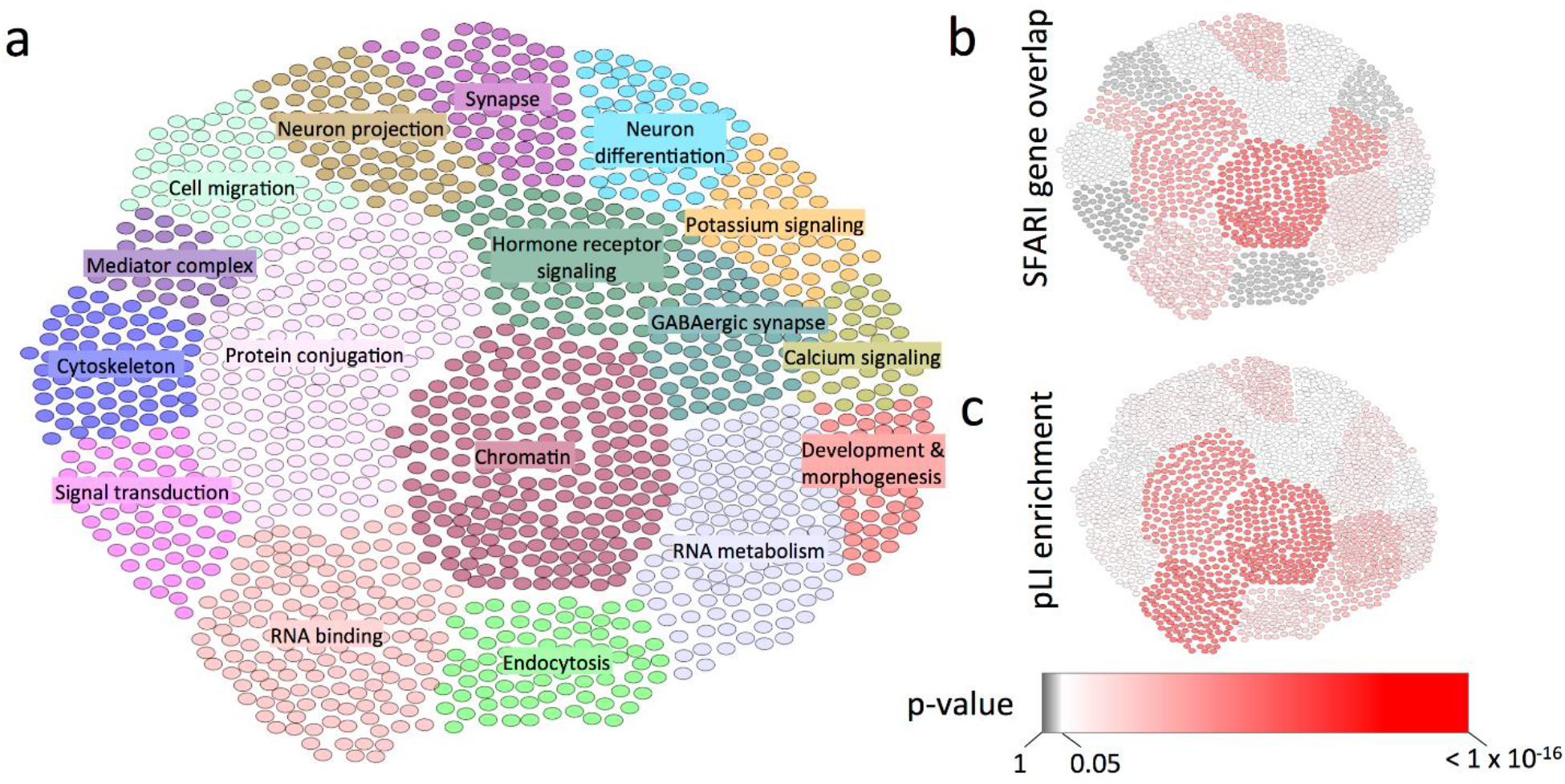
Clustering of top forecASD genes with ExAC pLI and SFARI high-confidence gene overlap enrichment analysis.. Greedy hierarchical optimization of the modularity score yielded 17 clusters consisting of 1452 forecASD genes (a). All clusters have several significantly enriched biological pathways, of which the top terms were overlaid in figure 6A. Clusters were tested for significance of overlap with the list of SFARI HC genes (b), and enrichment of haploinsufficiency genes (pLI > 0.5; c).

Lastly, forecASD genes were loaded into the STRING network and clustered using a greedy hierarchical approach which maximizes the modularity score. The resulting networks consisted of 17 clusters composed of 1452 genes. All clusters were found to be significantly enriched with numerous GO and KEGG pathways (Supplemental Table 3). Similarly, all clusters contained a significant enrichment of haploinsufficiency genes (pLI > 0.5), except for the small cluster of 31 genes with functions related to the mediator complex. Clusters were also tested for overlap with SFARI HC genes, of which 8 clusters failed to reach significance, suggesting groupings of genes currently missing from the known list of autism genes. Clusters lacking significant overlap includes those with functions: signal transduction, cytoskeleton, cell migration, neuron projection, steroid signaling, neuron differentiation, potassium signaling, development and morphogenesis. A marginally significant correlation was seen between a clusters enrichment for high pLI genes and its overlap with SFARI HC genes (Spearman’s r = 0.48, p-value = 0.053), further suggesting a bias in SFARI HC genes towards haploinsufficiency status.

## Discussion

We present forecASD, a machine learning approach that combines systems biology and genetic models into a single score that indexes the strength of evidence for a gene’s involvement in autism. This genome-wide score can be a useful prior, filter, or positive control in molecular studies of autism. It can also be used as a starting point to generate new hypotheses to investigate currently under-appreciated aspects of the molecular etiology of autism. In our tests of predictive performance and generalization, forecASD outperformed other systems biology and genetic approaches for autism gene prioritization.

Because it draws upon multiple approaches for identifying autism genes, forecASD is less biased than gene discovery based only on one form of data (e.g., genetic data). This is particularly important because current SFARI HC genes, which rely heavily on studies of *de novo* mutation, are strongly biased towards genes that are loss-of-function intolerant (Fig. 5A). While these haploinsufficient genes represent a sizable and important component of genetic risk for autism, this ascertainment bias has led to molecular “blind spots” that will not be resolved simply by sequencing more probands and identifying additional *de novo* mutations. For instance, pathways implicated preferentially by SFARI HC genes had significantly higher pLI, whereas pathways with lower pLI were under-represented (compared to forecASD-implicated pathways; OR=0.38, P=5.6×10^-7^). Furthermore, while SFARI HC pathways significantly predicted case/control expression-implicated pathways (Z=4.5, P=7.9×10^-6^, binomial model), only 3% of the deviance could be explained. In contrast, forecASD pathways explained an order of magnitude more deviance (31%, Z=12.1, P=1.5×10^-33^) when predicting expression-implicated pathways. When a model of dichotomous DE significance was fit that included terms from both SFARI HC and forecASD pathways, the SFARI HC term became redundant, and the forecASD-only model yielded a superior Bayesian information criterion (BIC; 518 for forecASD-only vs. 521 for full model and 720 for the SFARI HC-only model). When considering directionality, SFARI HC gene pathways were significantly depleted for ASD-upregulated pathways (OR=0.48, P=1.3×10^-6^), further illustrating the bias in SFARI HC genes. These results demonstrate that forecASD showed greater representation of low pLI and ASD-upregulated pathways, without sacrificing sensitivity to well-known ASD risk pathways where haploinsufficiency plays a dominant role.

In our analyses, we noted a trend that is a potential bridge between gene discovery studies based on DNA sequence variants and those based on differential expression. Specifically, pLI is significantly and negatively correlated with previously published frontal cortex differential expression (case/control) t-statistics(17) (beta=-0.13, t-statistic-4.3, P=1.9×10^-5^). This suggests that low-pLI genes are more likely to be up-regulated and high-pLI genes are more likely to be down-regulated in ASD cases (Fig. 5B). This is consistent with our observation of SFARI HC gene pathways (which have an ascertainment bias in favor of haploinsufficiency) being significantly under-represented in both low pLI and ASD-upregulated pathways. We further observed a significant interaction (*F*=54.1, P=3.9×10^-24^) between forecASD and pLI when explaining variation in ASD brain gene expression: forecASD genes (i.e., top decile) show the significant negative relationship between pLI and t-statistic (beta=-0.24, t=-2.6, P=0.009; Fig. 5D), while non-forecASD genes show no relationship (beta=0.004, t=0.1, P=0.91; Fig. 5C). Consequently, we propose that this pLI-expression relationship is a hallmark of robust ASD risk genes, and may be used as a criterion when identifying optimal thresholds in genome-wide scores like forecASD. Indeed, although initially chosen as a convenient but arbitrary threshold for identifying a discrete set of ASD candidate genes, the top decile proved to be the optimal split point for forecASD, maximizing the significance of the pLI/t-statistic relationship among candidate genes, while minimizing the same relationship in the remaining, non-candidate genes. Interestingly, when applying this approach to TADA FDR values, although TADA-implicated genes showed the expected pLI-expression relationship, no TADA threshold was able to eliminate the trend from non-candidate genes, suggesting lower sensitivity in identifying ASD risk genes compared to forecASD. Taken together, these analyses demonstrate that the reduced bias in forecASD contributes to increased sensitivity to autism risk pathways identified in gene expression studies (Fig. 4C,4D) as well as those implicated by genetic studies (Fig. 3).

Some pathways, although represented (but not necessarily enriched) in the current SFARI HC list, showed a substantial relative increase in enrichment when considering forecASD (Fig. 4A, Supplemental Table 2). This suggests that these pathways represent noted and plausible but still under-appreciated molecular themes in our understanding of autism. The pathway that underwent the largest relative increase in enrichment from SFARI HC to forecASD is Rho GTPase signaling (OR=2.2, P=4.8×10^-5^), which plays a critical role in cytoskeletal dynamics in neurodevelopment(24), including interactions with SHANK proteins and the formation and maturation of dendritic spines(25). As another example, although chromatin modification in general is a well-established theme in autism genetic risk, histone acetyltransferases showed relatively little representation in the SFARI HC list, but were significantly enriched in forecASD genes (OR=4.1, P=3.5×10^-9^). Histone acetylation was recently shown to be a pervasive genomic predictor of affected status in a large autism case/control postmortem brain study(26), underscoring the importance of this mechanism that is under-represented in established risk genes but that forecASD was sensitive to. As a final example of these under-appreciated molecular mechanisms, the circadian clock pathway was implicated by forecASD as an important source of risk for autism (OR=6.5, P=7.6×10^-13^). Sleep disturbances are a well-known and problematic comorbidity in autism, and molecular deficits in circadian regulation related to autism have been documented(27)^,(28),(29)^. Although literature support is available for these processes playing a role in autism, our results indicate that their current sparse representation in lists of accepted genetic risk factors is not representative of their importance in the disorder.

Other pathways were identified by forecASD as significantly enriched for autism risk, but were not represented at all among SFARI HC genes (Supplemental Table 2, Figure 4B). Consequently, we expect that new insights into the molecular basis of autism will come disproportionately from these pathways as their constituent genes are associated with autism. One gene set in particular, potassium channels, showed highly significant enrichment in forecASD genes (OR=4.1, P=7.2×10^-9^, N=35 genes) despite the absence of potassium channel genes among currently accepted autism risk genes. However, the literature shows support for a role for potassium channels in ASD risk(30)^,(31),(32),(33)^, and the pathway was enriched for differential regulation in a recently published brain gene expression study of autism (P=0.001, downregulated)(17). Notably, this pathway has a lower proportion of genes with pLI>0.9 (0.22) compared to SFARI HC gene-implicated pathways (median=0.47), potentially explaining its absence due to ascertainment bias. Overall, pathways that demonstrated forecASD-specific excess enrichment showed a significant agreement with pathway enrichment from independent case/control brain gene expression studies (OR=28.8; P=2.9×10^-48^), and were more likely to support pathways that were up-regulated in the gene expression data (OR=2.1, P=1.27×10^-6^, Fig. 4C) compared to pathways implicated by SFARI HC.

To group forecASD genes into distinct functional categories, we performed iterative clustering and identified a total of 17 clusters enriched for specific functional annotations. While nearly all clusters showed significant enrichment for haploinsufficiency genes, many lacked a significant overlap with SFARI HC genes, after Bonferroni correction. Similar to conclusions reached above, we found an entire cluster enriched for Potassium signaling (P=1.8×10^-49^) which lacked significant overlap with SFARI HC genes. In addition to this cluster, there were also seven others lacking significant overlap with SFARI HC genes. Notable examples include clusters related to cell migration (P=6.0×10^-11^) and endocytosis (P=2.7×10^-21^). These pathways have more recently been explored in their ability to regulate brain connectivity(34) and postsynaptic organization(35), respectively. In agreement with the proposed haploinsufficiency bias of autism gene discovery, we observed a marginally significant relationship between cluster pLI enrichment and SFARI HC gene overlap (Spearman’s rho: 0.48; P=0.053).

During the development of forecASD, another ASD gene prediction method was published(36), ASD-FRN, which utilizes a brain-specific functional network. We evaluated this method using the performance benchmarks presented in Figure 3, and found it to have performance comparable to DAMAGES and Krishnan, et al. (Supplemental Figure 1 and Supplemental Figure 2), but in none of these tests did it surpass the performance observed using forecASD as an ASD gene predictor. We also added ASD-FRN to the ensemble that comprises forecASD, but we did not observe a significant increase in predictive performance, suggesting that the latent information in ASD-FRN is already accounted for in the forecASD ensemble as presented here.

Our use of TADA to impart genetic association information to the forecASD ensemble is unique among the ASD gene prediction approaches we used as benchmarks. However, this raises concerns about the potential for circularity: TADA is emerging as the most popular way to compute and update gene-wise genetic association statistics for ASD studies, and previous TADA scores are strongly correlated with updated TADA scores. Furthermore, TADA scores are among the most important predictive features in the forecASD ensemble (Fig. 2). Consequently, it would be concerning if forecASD’s ability to predict new ASD genes was due entirely to the inclusion of previously published TADA scores. To examine this possibility, we fit bivariate logistic regression models, always including TADA rankings as a covariate, with Krishnan, DAMAGES, ASD-FRN, or forecASD rankings as the predictor of membership among SFARI Gene score 3 (Supplemental Figure 2; score 3 genes were not used in training forecASD, but are nevertheless assumed to be enriched for true ASD genes). All other genes listed in the SFARI Gene database were removed from the analysis, and the remainder of the genome was considered the negative class. Strong association with score 3 membership (as measured by the regression coefficient and Z-score) is a desirable trait for an ASD gene prediction method, and forecASD proved to be the most strongly associated of the tested methods, even after correcting for the information imparted by TADA rankings (Z=11.9, P=8.9×10^-33^).

Rather than simply parroting gene prioritization according to TADA, forecASD purifies TADA’s signal by imposing biological plausibility through the inclusion of other predictive features, such as network and expression data, as well as previously published machine learning approaches to ASD gene prediction. To illustrate this filtering effect that forecASD has on TADA rankings, we performed a GSEA on the differences in gene ranks between forecASD and TADA. This analysis highlights pathways and processes favored by forecASD, vs. those favored by TADA (Supplemental Figure 3). Pathways where there is consensus fall in the middle of the difference-in-ranks distribution, and consequently are not highlighted as enriched or depleted in this analysis. In this analysis, forecASD prioritized genes involved in well-established neurodevelopmental pathways, such as signaling by receptor tyrosine kinases, axon guidance, Rho GTPase signaling, and chromatin modification. Conversely, TADA prioritized genes belonging to large gene families, such as olfactory receptors, ribosomal proteins, and defensins. These kinds of genes are routinely manually removed from consideration even in the presence of statistical support, because they often lack biological plausibility. forecASD accomplishes this same task, but in a more automated, data-driven, and objective way.

One bias in forecASD uncovered by the above analysis, and that is intuitive based on forecASD’s reliance on functional data, is that forecASD tends to de-prioritize genes that are poorly annotated or lack strong signals in expression or other functional data sets (in the above GSEA, forecASD-preferred genes were depleted for “Uncategorized” at P=6.8×10^-9^). On one hand, this limits forecASD’s ability to make entirely unexpected discoveries among poorly characterized genes. On the other hand, the probability of highly important genes completely evading decades of neuroscience investigation is diminishing, and penalizing these “unassuming” genes may be justifiable. Nevertheless, the value of new discovery should not be discounted, and this will be an area of active investigation in the ongoing development of forecASD.

As a final proof of principle for one of forecASD’s intended uses, we investigated genes nominated as new ASD genes by a currently unpublished large-scale gene discovery study(37). The data underlying this study was not available to us during the development of forecASD, so this represents a true test of forecASD’s generalization and interpretive value. This study nominated *BTRC*, *C16orf13*, *CCSER1*, *CMPK2*, *DDX3X*, *FAM98C*, *GRIA1*, *MLANA*, *MYO5A*, *PCM1*, *PRKAR1B*, *RAPGEF4*, *SMURF1*, *TMEM39B*, *TSPAN4*, and *UIMC1* as newly discovered ASD genes at a TADA FDR of 0.2, meaning that of these 16 genes, 2-3 are expected to be false positives. Overall, the nominated genes are strongly enriched for elevated forecASD scores (P=4.8×10^-8^, Wilcoxon test), and 12 of the 16 genes scored in the top decile of forecASD. This result strengthens both the case for forecASD as an effective predictor of ASD risk genes, and the findings of this gene discovery study. However, *FAM98C* and *CCSER1* have very low forecASD scores (0.022 and 0.024, respectively), indicating little support in network and functional data. *UIMC1* and *C16orf13* have slightly higher scores (0.210 and 0.346, respectively), but are still below the top-decile threshold we have used in this study (0.374). Consequently, because of weak support beyond the genetic association, these four genes are candidates for being the false positives, and should require additional experimental data supporting a role in autism before they are considered true ASD risk genes.

For the foreseeable future, traditional gene discovery studies will continue to add to the list of bona fide ASD risk genes. Eventually, as sample sizes saturate and gene discovery decelerates, the field will be faced with the challenge of developing new and useful applications of this acquired knowledge. By combining new and previously published predictors into a high-performance ensemble classifier, forecASD provides a glimpse of that future and gives an opportunity right now to begin thinking about what we would do with a definitive list of autism genes.

## Conclusions

We introduce a new model, forecASD, for prioritizing autism-associated risk genes. We show that forecASD has significantly improved sensitivity and specificity compared to previous gene-level predictors, including the purely genetic-based approach TADA. We demonstrate forecASD’s usefulness as an autism risk-gene discovery post-hoc filter through its ability to prioritize 12 of 16 genes implicated at FDR=0.2 by the latest ASD gene discovery study. Using forecASD prioritized risk-genes, we highlight which molecular pathways are currently under-represented in the autism literature and likely represent under-appreciated biological mechanisms of autism.

## List of abbreviations

LGD: likely gene disrupting
DNM: de novo mutation
TADA: Transmission And De novo Association
DE: differential expression
GSEA: gene set enrichment analysis

## Declarations

### Ethics approval and consent to participate

All human genetic data used in this study was accessed in a deidentified manner from associated sequencing consortia with their approval. Due to the method of access, this study is not formally considered human subjects research and therefore not subject to associated restrictions, as outlined by the National Institutes of Health.

### Consent for publication

Not applicable.

### Availability of data and material

All data and code used to generate the forecASD model is available for download from the repository associated with this project, https://github.com/LeoBman/forecASD.

### Authors’ contributions

LB contributed to design, testing, and implementation of forecASD model and was a major contributor in writing the manuscript. TK contributed to analysis of forecASD model results and contributed to github implementation of forecASD. JM contributed to design, testing, and implementation of forecASD model and was a major contributor in writing the manuscript. All authors read and approved the final manuscript.

## Acknowledgements

We are grateful to all of the families in SPARK, the SPARK clinical sites and SPARK staff. We appreciate obtaining access to exome sequencing and phenotypic data on SFARI Base. Approved researchers can obtain the SPARK population dataset described in this study by applying at https://base.sfari.org.

## References

1. Rosenberg RE, Law JK, Yenokyan G, McGready J, Kaufmann WE, Law PA. Characteristics and concordance of autism spectrum disorders among 277 twin pairs. Arch Pediatr Adolesc Med. 2009;163(10):907–14.

2. Colvert E, Tick B, McEwen F, Stewart C, Curran SR, Woodhouse E, et al. Heritability of Autism Spectrum Disorder in a UK Population-Based Twin Sample. JAMA Psychiatry. 2015;72(5):415–23.

3. De Rubeis S, He X, Goldberg AP, Poultney CS, Samocha K, Cicek AE, et al. Synaptic, transcriptional and chromatin genes disrupted in autism. Nature. 2014;515(7526):209–15.

4. Abrahams BS, Arking DE, Campbell DB, Mefford HC, Morrow EM, Weiss LA, et al. SFARI Gene 2.0: a community-driven knowledgebase for the autism spectrum disorders (ASDs). Mol Autism. 2013;4(1):36.

5. Liu L, Lei J, Sanders SJ, Willsey AJ, Kou Y, Cicek AE, et al. DAWN: a framework to identify autism genes and subnetworks using gene expression and genetics. Mol Autism. 2014;5(1):22.

6. Krishnan A, Zhang R, Yao V, Theesfeld CL, Wong AK, Tadych A, et al. Genome-wide prediction and functional characterization of the genetic basis of autism spectrum disorder. Nat Neurosci. 2016;19(11):1454–62.

7. Zhang C, Shen Y. A Cell Type-Specific Expression Signature Predicts Haploinsufficient Autism-Susceptibility Genes. Hum Mutat. 2017;38(2):204–15.

8. RK CY, Merico D, Bookman M, J LH, Thiruvahindrapuram B, Patel RV, et al. Whole genome sequencing resource identifies 18 new candidate genes for autism spectrum disorder. Nat Neurosci. 2017;20(4):602–11.

9. SCEa, Consortium S. SPARK: A US Cohort of 50,000 Families to Accelerate Autism Research. Neuron. 2018;97(3):488–93.

10. Sunkin SM, Ng L, Lau C, Dolbeare T, Gilbert TL, Thompson CL, et al. Allen Brain Atlas: an integrated spatio-temporal portal for exploring the central nervous system. Nucleic Acids Res. 2013;41(Database issue):D996–D1008.

11. von Mering C, Huynen M, Jaeggi D, Schmidt S, Bork P, Snel B. STRING: a database of predicted functional associations between proteins. Nucleic Acids Res. 2003;31(1):258–61.

12. Sanders SJ, He X, Willsey AJ, Ercan-Sencicek AG, Samocha KE, Cicek AE, et al. Insights into Autism Spectrum Disorder Genomic Architecture and Biology from 71 Risk Loci. Neuron. 2015;87(6):1215–33.

13. Fabregat A, Jupe S, Matthews L, Sidiropoulos K, Gillespie M, Garapati P, et al. The Reactome Pathway Knowledgebase. Nucleic Acids Res. 2018;46(D1):D649–D55.

14. Mi H, Lazareva-Ulitsky B, Loo R, Kejariwal A, Vandergriff J, Rabkin S, et al. The PANTHER database of protein families, subfamilies, functions and pathways. Nucleic Acids Res. 2005;33(Database issue):D284–8.

15. R Development Core Team. R: A language and environment for statistical computing. Vienna, Austria: R Foundation for Statistical Computing; 2008.

16. Lek M, Karczewski KJ, Minikel EV, Samocha KE, Banks E, Fennell T, et al. Analysis of protein-coding genetic variation in 60,706 humans. Nature. 2016;536(7616):285–91.

17. Gandal MJ, Haney JR, Parikshak NN, Leppa V, Ramaswami G, Hartl C, et al. Shared molecular neuropathology across major psychiatric disorders parallels polygenic overlap. Science. 2018;359(6376):693–7.

18. Szklarczyk D, Morris JH, Cook H, Kuhn M, Wyder S, Simonovic M, et al. The STRING database in 2017: quality-controlled protein-protein association networks, made broadly accessible. Nucleic Acids Res. 2017;45(D1):D362–D8.

19. Sugathan A, Biagioli M, Golzio C, Erdin S, Blumenthal I, Manavalan P, et al. CHD8 regulates neurodevelopmental pathways associated with autism spectrum disorder in neural progenitors. Proc Natl Acad Sci U S A. 2014;111(42):E4468–77.

20. Darnell JC, Van Driesche SJ, Zhang C, Hung KY, Mele A, Fraser CE, et al. FMRP stalls ribosomal translocation on mRNAs linked to synaptic function and autism. Cell. 2011;146(2):247–61.

21. Csardi G NT. The igraph software package for complex network research. InterJournal 2006.

22. Newman ME. Modularity and community structure in networks. Proc Natl Acad Sci U S A. 2006;103(23):8577–82.

23. Shannon P, Markiel A, Ozier O, Baliga NS, Wang JT, Ramage D, et al. Cytoscape: a software environment for integrated models of biomolecular interaction networks. Genome Res. 2003;13(11):2498–504.

24. Reichova A, Zatkova M, Bacova Z, Bakos J. Abnormalities in interactions of Rho GTPases with scaffolding proteins contribute to neurodevelopmental disorders. J Neurosci Res. 2018;96(5):781–8.

25. Martin-Vilchez S, Whitmore L, Asmussen H, Zareno J, Horwitz R, Newell-Litwa K. RhoGTPase Regulators Orchestrate Distinct Stages of Synaptic Development. PLoS One. 2017;12(1):e0170464.

26. Sun W, Poschmann J, Cruz-Herrera Del Rosario R, Parikshak NN, Hajan HS, Kumar V, et al. Histone Acetylome-wide Association Study of Autism Spectrum Disorder. Cell. 2016;167(5):1385–97 e11.

27. Lipton JO, Boyle LM, Yuan ED, Hochstrasser KJ, Chifamba FF, Nathan A, et al. Aberrant Proteostasis of BMAL1 Underlies Circadian Abnormalities in a Paradigmatic mTOR-opathy. Cell Rep. 2017;20(4):868–80.

28. Monyak RE, Emerson D, Schoenfeld BP, Zheng X, Chambers DB, Rosenfelt C, et al. Insulin signaling misregulation underlies circadian and cognitive deficits in a Drosophila fragile X model. Mol Psychiatry. 2017;22(8):1140–8.

29. Kozlov SV, Bogenpohl JW, Howell MP, Wevrick R, Panda S, Hogenesch JB, et al. The imprinted gene Magel2 regulates normal circadian output. Nat Genet. 2007;39(10):1266–72.

30. Guglielmi L, Servettini I, Caramia M, Catacuzzeno L, Franciolini F, D’Adamo MC, et al. Update on the implication of potassium channels in autism: K(+) channelautism spectrum disorder. Front Cell Neurosci. 2015;9:34.

31. Deng PY, Klyachko VA. Genetic upregulation of BK channel activity normalizes multiple synaptic and circuit defects in a mouse model of fragile X syndrome. J Physiol. 2016;594(1):83–97.

32. Lee H, Lin MC, Kornblum HI, Papazian DM, Nelson SF. Exome sequencing identifies de novo gain of function missense mutation in KCND2 in identical twins with autism and seizures that slows potassium channel inactivation. Hum Mol Genet. 2014;23(13):3481–9.

33. Sicca F, Ambrosini E, Marchese M, Sforna L, Servettini I, Valvo G, et al. Gain-of-function defects of astrocytic Kir4.1 channels in children with autism spectrum disorders and epilepsy. Sci Rep. 2016;6:34325.

34. Reiner O, Karzbrun E, Kshirsagar A, Kaibuchi K. Regulation of neuronal migration, an emerging topic in autism spectrum disorders. J Neurochem. 2016;136(3):440–56.

35. Loebrich S. The role of F-actin in modulating Clathrin-mediated endocytosis: Lessons from neurons in health and neuropsychiatric disorder. Commun Integr Biol. 2014;7:e28740.

36. Duda M, Zhang H, Li HD, Wall DP, Burmeister M, Guan Y. Brain-specific functional relationship networks inform autism spectrum disorder gene prediction. Transl Psychiatry. 2018;8(1):56.

37. Ruzzo EK, Perez-Cano L, Jung J-Y, Wang L-k, Kashef-Haghighi D, Hartl C, et al. Whole genome sequencing in multiplex families reveals novel inherited and de novo genetic risk in autism. bioRxiv. 2018.

